# An eutherian intronic sequence gave rise to a major satellite DNA in Platyrrhini

**DOI:** 10.1101/213868

**Authors:** Mirela Pelizaro Valeri, Guilherme Borges Dias, Valéria do Socorro Pereira, Gustavo Campos Silva Kuhn, Marta Svartman

**Affiliations:** Laboratório de Citogenômica Evolutiva, Departamento de Biologia Geral, Instituto de Ciências Biológicas, Universidade Federal de Minas Gerais; Belo Horizonte, MG, Brazil; Fundação Zoo-Botânica de Belo Horizonte, Belo Horizonte, MG, Brazil

**Keywords:** Repetitive DNA, Satellite DNA origin, Platyrrhini

## Abstract

Satellite DNAs (satDNAs) are major components of eukaryote genomes. However, because of their quick divergence, the evolutionary origin of a given satDNA family can rarely be determined. Herein we took advantage of available primate sequenced genomes to determine the origin of the CapA satDNA (~1,500 bp long monomers), first described in *Sapajus apella*. We show that CapA is an abundant satDNA in Platyrrhini, whereas in the genomes of most eutherian mammals, including humans, this sequence is present only as a single copy located within a large intron of the NOS1AP (*nitric oxid synthase 1 adaptor protein*) Gene. Our data suggest that this intronic CapA-like sequence gave rise to the CapA satDNA and we discuss possible mechanisms implicated in this event. This is the first report of a single copy intronic sequence giving origin to a satDNA that reaches up to 100,000 copies in some genomes.

---

Eukaryote genomes are replete with repetitive DNA sequences amongst which satellite DNAs (satDNAs) are usually prominent components [1]. These tandem repeats (TRs) are commonly found as very long arrays located in heterochromatic regions of chromosomes, such as pericentric and centromeric heterochromatin, and subtelomeric regions [2]. Because satDNAs are fast-evolving, their origin is often hard to determine [1, 2]. The most recurrent evolutionary path suggested for satDNA origin is the tandem amplification of internal segments of transposable elements (TEs), with many cases described so far in animals and plants [3].

The study of primate genomes is paramount for medical genetics and comparative evolutionary studies of the human genome. Nevertheless, as for most eukaryotes, the satDNA fraction of these genomes is often overlooked. Herein, we took advantage of the available set of primate sequenced genomes to study the evolution of a satDNA called CapA. This satDNA was first described in the New World monkey (NWM) *Sapajus apella* (at the time classified as *Cebus apella,* Cebidae, Platyrrhini). It has large monomers of ~1,500 bp, and DNA-DNA hybridization experiments revealed its presence in other NWMs [4, 5]. However, because of the paucity of DNA sequence data available at the time of its description, CapA origin and evolution remained elusive.

A similarity search against the NCBI nucleotide collection using BLASTn and the CapA sequence described in Malfoy et al. [4] as a query returned highly similar sequences from several mammals (Supplementary Table 1). Interestingly, one of these sequences (77% coverage with 79% identity) was annotated as the *Homo sapiens nitric oxid synthase 1 adaptor protein* (NOS1AP), located at the proximal region of the long arm on chromosome 1 (1q23.3). A closer inspection of this gene revealed that the region with similarity to the deposited CapA monomer is ~1,180 bp long and is situated in the second intron of NOS1AP. Subsequent BLAT searches using this intronic CapA segment against the whole human genome (hg38) retrieved no additional matches, indicating that this CapA-like sequence is not repetitive in humans.

We then proceeded to use this human CapA-like intronic sequence as a query against all vertebrate genomes available in NCBI. This search retrieved similar sequences mostly as single matches in each genome with CapA-like sequences being present in most eutherians. Data from a few species with partial chromosome assembly and chromosome painting data with human probes confirm that the CapA-like sequence remains in the same locus, probably also associated with the NOS1AP gene (Supplementary Table 2). However, some eutherian clades appear to lack this CapA-like sequence, most notably the Chiroptera, some Eulipotyphla and some Rodentia (Supplementary Table 2, Figure 1). No CapA-like sequences were found in Marsupialia or Monotremata, which are the sister clades to Eutheria. These findings suggest that a single CapA-like sequence was present in the ancestor of eutherians and that it was likely the precursor sequence that gave rise to the CapA satellite DNA (Figure 1).

**Figure 1.**
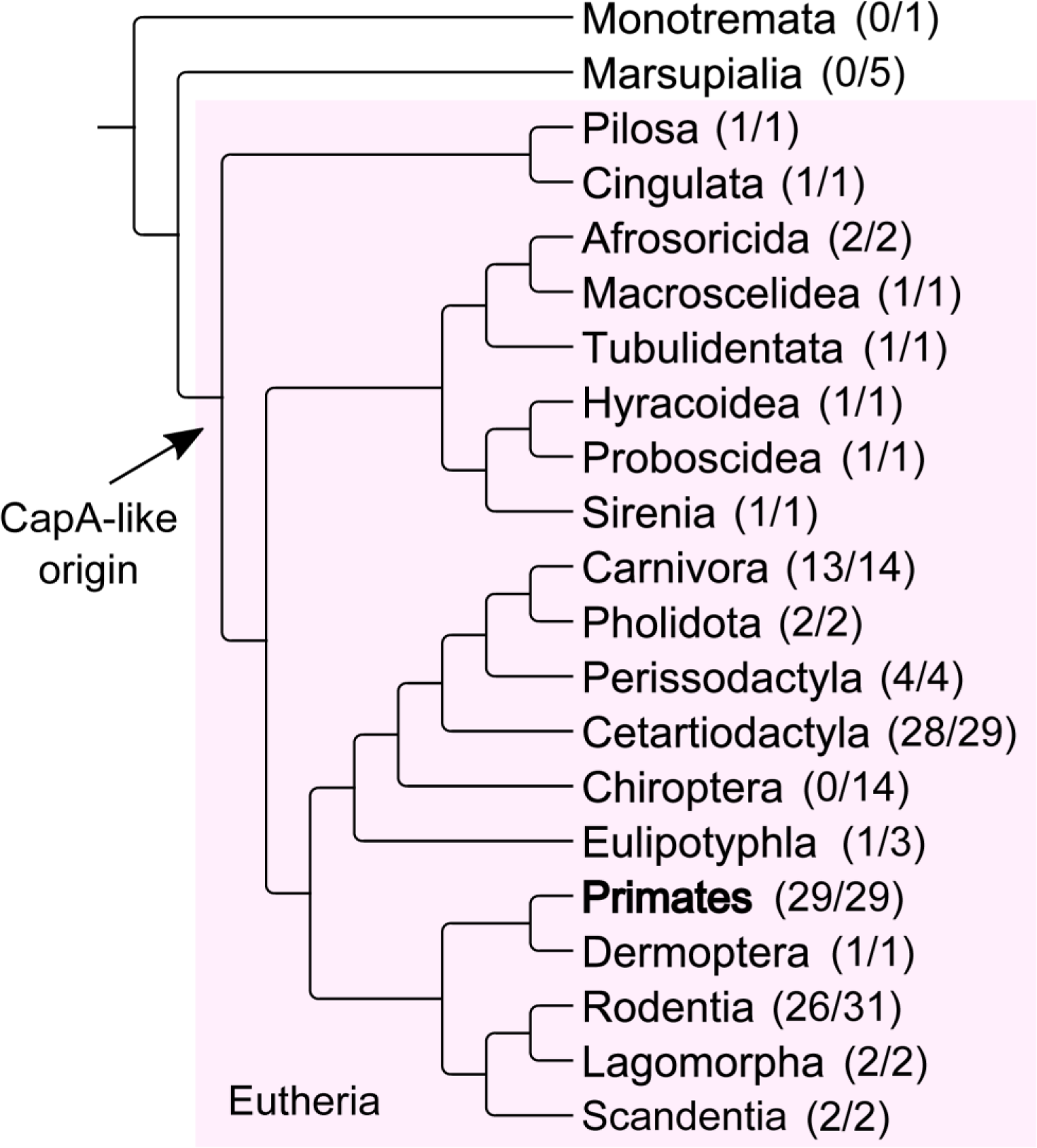
Diagrammatic representation of the mammalian phylogeny adapted from Foley et al. [14]. The CapA precursor probably predates the eutherian radiation (arrow) and was likely lost in a few clades. Numbers in parentheses indicate the genomes of species in which CapA was found and the total number of species analyzed. Within primates (bold) CapA was found as a satDNA only in Platyrrhini.

The only genomes in which we found multiple copies of CapA were those of NWMs from the family Cebidae, in which some contigs revealed the presence of a few CapA TRs (Supplementary Table 3). However, because the assembly of TRs is usually incomplete, especially for TRs with monomers larger than read sizes (as in the case of CapA), we also used raw Illumina data to estimate CapA abundance in available NWMs genomes (Supplementary Table 4). Strikingly, this approach revealed that CapA is an abundant tandem repeat in *Cebus capucinus* (genome proportion of 4.21%), *S. boliviensis* (1.48%) and *A. nancymaae* (0.27%) (Supplementary Table 4). In fact, CapA divergence landscapes are very similar across the three Cebidae genera, despite the significant differences in its abundance (Supplementary Figure 1). A slightly higher overall divergence was detected in *S*. *boliviensis* (19.67%), compared to 17.8 and 18.46% in *A*. *nancymaae* and *C*. *capucinus*, respectively (Supplementary Figure 1). In *Callithrix jacchus,* we found CapA in low copy numbers but also tandemly arranged (Supplementary Table 3). Interestingly, even in the genomes where CapA achieved high abundance, the intronic ancestral locus persisted as an intact CapA-like monomer, with the exception of *Callithrix jacchus*. In this species the intronic CapA on chromosome 1 suffered a rearrangement involving a small duplication and the insertion of an unrelated sequence of ~300 bp (Supplementary Figure 2).

All the currently available genomes of NWMs belong to species of the Cebidae family, preventing us from assessing the amplification status of CapA in the other two NWMs families, Atelidae and Pitheciidae. To expand our analysis to species without sequence data, we assessed CapA abundance by performing fluorescent in situ hybridization (FISH) with the human intronic CapA-like sequence as a probe onto cells of several NWMs (Table 1). We detected signs of CapA expansion in members of the three Platyrrhini families (Figure 2). Within Cebidae, CapA was present in high abundance in *Sapajus xanthosternos*, *Saimiri boliviensis* and *Aotus infulatus* and was not detected in representatives of Callithrichinae (a Cebidae subfamily). CapA also occurred as a high copy sequence in *Alouatta guariba*, *Lagothrix lagotricha* and *Brachyteles hypoxanthus*, of the family Atelidae, in which CapA was less abundant in *A. guariba* than in *L. lagotricha* and *B. hypoxanthus*. In Pitheciidae, *Chiropotes satanas* and *Pithecia irrorata* also presented signs of CapA expansion, whereas *Callicebus nigrifrons* did not. CapA was found in only one small acrocentric chromosome pair of *P. irrorata* and was very abundant in *C. satanas*. In all the species in which CapA was abundant, it was associated with heterochromatin. In the species that did not display visible blocks of CapA it is possible that it occurs in very low copy numbers or that the sequence has diverged considerably, preventing its detection by FISH.

**Figure 2.**
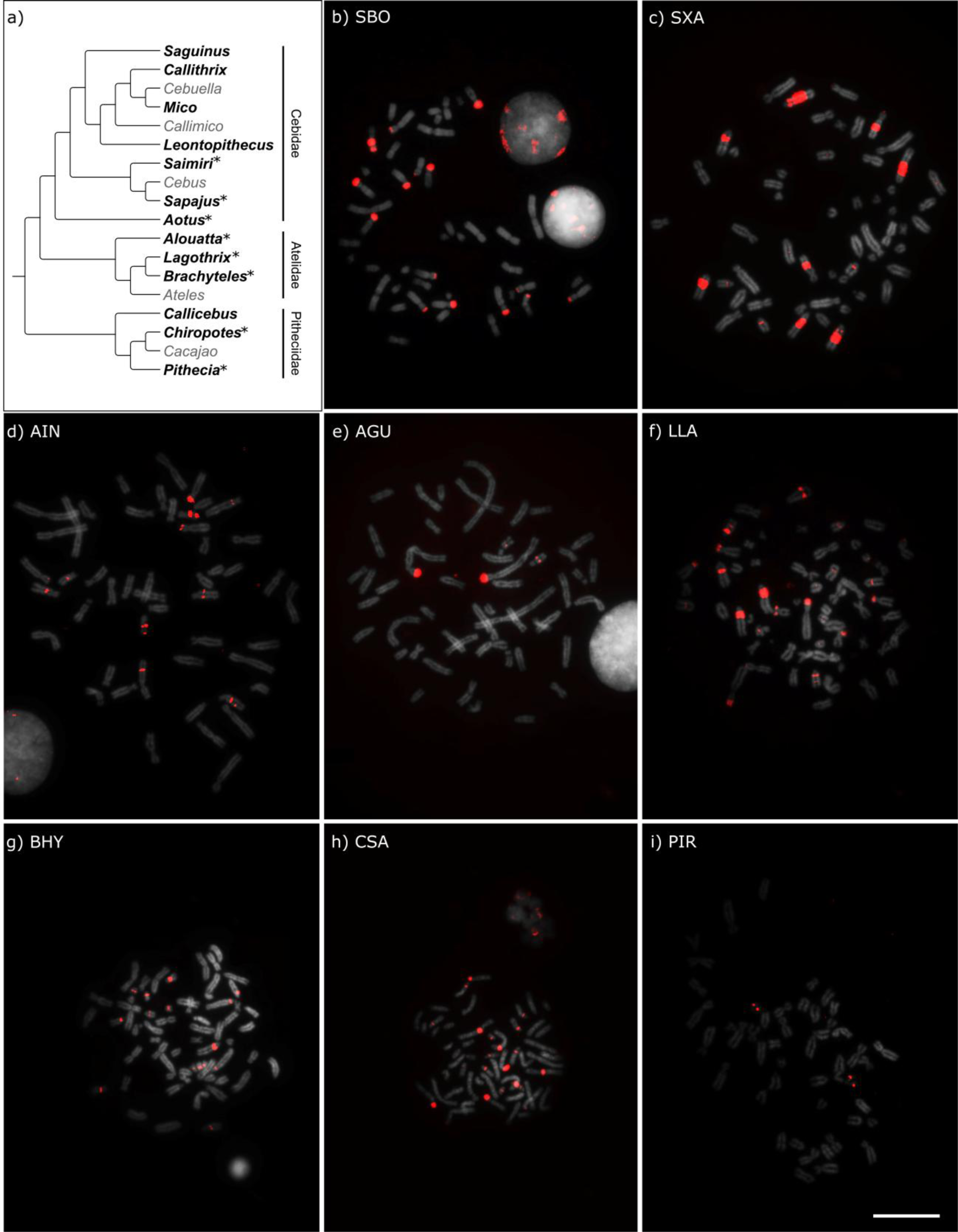
a) Distribution of CapA among Platyrrhini families. Diagrammatic representation of the Platyrrhini phylogeny adapted from Schneider & Sampaio [5]. The genera with representatives analyzed are shown in bold and the asterisk indicates CapA amplification. Fluorescent *in situ* hybridization with CapA in metaphases of: b) *Saimiri boliviensis* (SBO), c) *Sapajus xanthosternos* (SXA), d) *Aotus infulatus* (AIN), e) *Alouatta guariba* (AGU), f) *Lagothrix lagotricha* (LLA), g) *Brachyteles hypoxanthus* (BHY), h) *Chiropotes satanas* (CSA) and i) *Pithecia irrorata* (PIR). Bar = 10μm.

**Table 1.** Species in which FISH with the human intronic CapA-like was performed.

| Species | Family | Procedence |
| --- | --- | --- |
| *Saguinus imperator* (SIM) | Cebidae | Fundação Zoo-Botânica de Belo Horizonte |
| *Callithrix penicillata* (CPE) | Cebidae | Universidade Federal de Minas Gerais |
| *Mico argentata* (MAR) | Cebidae | Universidade de São Paulo |
| *Leontopithecus rosalia* (LRO) | Cebidae | Fundação Zoo-Botânica de Belo Horizonte |
| *Sapajus xanthosternos* (SXA) | Cebidae | Fundação Zoo-Botânica de Belo Horizonte |
| *Saimiri boliviensis* (SBO) | Cebidae | University of Florence |
| *Aotus infulatus* (AIN) | Cebidae | Fundação Zoo-Botânica de Belo Horizonte |
| *Alouatta guariba* (AGU) | Atelidae | Fundação Zoo-Botânica de Belo Horizonte |
| *Lagothrix lagotricha* (LLA) | Atelidae | Fundação Zoo-Botânica de Belo Horizonte |
| *Brachyteles hypoxanthus* (BHY) | Atelidae | Fundação Zoo-Botânica de Belo Horizonte |
| *Callicebus nigrifrons* (CNI) | Pitheciidae | Fundação Zoo-Botânica de Belo Horizonte |
| *Chiropotes satanas* (CSA) | Pitheciidae | Fundação Zoo-Botânica de Belo Horizonte |
| *Pithecia irrorata* (PIR) | Pitheciidae | Fundação Zoo-Botânica de Belo Horizonte |

The data presented herein suggest that CapA suffered an expansion within Platyrrhini, less than ~25 million years ago (Mya), before the NWMs families first diverged [6]. Assuming that CapA expansion predates the divergence of the three NWMs families, this satDNA would have been independently lost in some taxa, notably the Callithrichinae and *C. nigrifrons*. A second less parsimonious hypothesis would be that CapA has become a satDNA independently multiple times, as only some species within each family showed CapA expansion (Figure 2). To investigate the alternative possibilities more NWMs species need to be analyzed at the cytogenetic and genomic levels.

Herein we found that the large CapA satDNA present in Platyrrhini has originated from an intronic single copy sequence that is still present in most eutherians. Nevertheless, it is difficult to reconstruct the steps of CapA amplification. One possibility is that CapA arose through segmental duplications. In fact, segmental duplications have already been evoked to explain the hyperexpansion of sequences in primate genomes [7]. The single copy intronic precursor sequence is still present in the putative ancestral locus of CapA. Therefore, prior to amplification, this CapA sequence would have had to undergo transposition to another genomic region. Transposable elements (TEs), such as L1 retrotransposons (long interspersed nucleotide element–1), may have also participated in the process through transduction events. L1 is associated with the indirect spread of other retrotranscripts, but it can also carry non-L1 DNA that are flanking L1 3’ ends sequences to new genomic locations [8,9].

Duplicative transposition followed by expansion of particular euchromatic segments have been described in the pericentromeric regions of human and in the subterminal ends (generally heterochromatic) of great apes chromosomes [7,10]. CapA duplication and expansion in NWMs may be explained by a similar mechanism, in which transposition of the CapA intronic segment to heterochromatic regions in the ancestral Platyrrhini genome followed by hyperexpansion through unequal crossing over would have given rise to the CapA satDNA in some species. Although we favor this segmental duplication hypothesis, the incompleteness of current NWMs genome assemblies prevents its scrutiny to exhaustion.

In conclusion, we characterized CapA, a satDNA formed less than 25 Mya, with ~1,500bp monomers present in species of the three Platyrrhini families. In Cebidae, with the exception of Callitrichines, CapA abundance ranges from 0.27 to 5% of the genome. The CapA-like ancestral sequence is present in most eutherians, most likely embedded in the second intron of the *nitric oxide synthase 1 adaptor protein* (NOS1AP) gene, such as in *Homo sapiens*. One hypothesis for the CapA expansion in NWMs is through duplicative transposition followed by expansion through unequal crossing over. To the best of our knowledge, this is the first report of a single copy intronic sequence giving origin to a satDNA, with as much as 100,000 copies in some genomes.

## Material and Methods

We searched for sequences similar to CapA in the non-redundant nucleotide collection of GenBank using the CapA monomer described in Fanning et al. [5] as query and the BLASTn tool [11]. This search returned hits from several mammals, particularly NWMs, and included a hit in the intron of the *nitric oxide synthase 1 adaptor protein* (NOS1AP) gene from *Homo sapiens* (Supplementary Table 1). After checking the human reference genome (hg38) at UCSC using BLAT, we confirmed that this CapA-like sequence exists at a single locus, inside the NOS1AP gene on chromosome 1. We then used this human intronic sequence as query in BLASTn searches against all vertebrate assembled genomes available at NCBI. A hit was included when query cover was ≥30% and e-value ≤ e^−5^. Only Eutheria displayed CapA-like sequences. Number of hits with the CapA-like sequence, query cover, identity and E-value of each search are available in Supplementary table 2.

In the genomes where multiple hits of CapA were found, we used raw Illumina reads and RepeatMasker [12] to estimate CapA’s abundance. We included data from an Old World monkey (*Chlorocebus aethiops*) and a great ape (*Homo sapiens*) as negative controls. All sequence reads used in this step were obtained from the Short Read Archive at NCBI (available at http://www.ncbi.nlm.nih.gov/sra/), with accession numbers as follows: *Aotus nancymaae* SRR1692991, *Callithrix jacchus* SRR1746970, *Cebus capucinus imitator* SRR3144006, *Saimiri boliviensis* SRR317821, *Chlorocebus aethiops sabaeus* SRR5251202, and *Homo sapiens* ERR016352. Sequence reads were downloaded in fastq format using the software SRAToolkit (available at https://github.com/ncbi/sra-tools) and random samples of two million reads ranging from 101 to 125 bp were produced using the software seqtk (available at https://github.com/lh3/seqtk/). To determinate the fraction of the reads similar to CapA, RepeatMasker was used in the following setup: sensitive mode, without searching for low complexity or bacterial insertion sequences, using wublast as search engine and a custom library containing the NOS1AP intronic CapA-like sequence. We used the alignment files generated by RepeatMasker to calculated Kimura distances of CapA fragments against the ancestral sequence (NOS1AP intronic CapA-like) using the utility script calcDivergenceFromAlign.pl from the RepeatMasker package. Results were then imported to RStudio and plotted.

To investigate CapA abundance in species for which no genome data were available, we performed fluorescent in situ hybridization (FISH) with the intronic CapA probe. This probe was obtained after PCR amplification of human DNA with primers flanking the CapA-like intronic sequence from NOS1AP (CapA-F: ACTTCCTCACTGACCTGTCTT; CapA-R: GGGCTGATGCTTAATGTAGCA). The PCR products were purified, cloned and sequenced to ensure specificity (accession number: MG264524). Chromosome spreads of several NWMs, mostly with unknown geographic origin, were obtained from fibroblast or lymphocyte cultures (Table 1). FISH was performed with 200 ng of biotin labeled probes, following Araújo et al. [13].

## Authors’ contributions

MPV and GBD carried out the bioinformatics analyses and cytogenetic and molecular lab work, participated in data analysis and in the design of the study and drafting of the manuscript; VSP collected samples; GBD, GCSK and MS conceived and coordinated the study, and helped drafting the manuscript. All authors gave final approval for publication.

## Acknowledgements

The authors thank the following people for their assistance in obtaining samples for this study: Dr. Yatiyo Yonenaga-Yassuda and MSc Camila do Nascimento Moreira, Universidade de São Paulo; Dr. Roscoe Stanyon, University of Florence; Prof. Alan Lane de Melo, Universidade Federal de Minas Gerais.

